# A highly contiguous genome assembly of a major forest pest, the Eurasian spruce bark beetle *Ips typographus*

**DOI:** 10.1101/2020.11.28.401976

**Authors:** Daniel Powell, Ewald Groβe-Wilde, Paal Krokene, Amit Roy, Amrita Chakraborty, Christer Löfstedt, Heiko Vogel, Martin N Andersson, Fredrik Schlyter

**Affiliations:** Czech University of Life Sciences Prague, Faculty of Forestry and Wood Sciences, Excellent Team for Mitigation (ETM), Kamýcká 129, CZ – 165 00 Praha 6 – Suchdol, Czech Republic; Department of Biology, Lund University, SE-223 62 Lund, Sweden; Division of Biotechnology and Plant Health, Norwegian Institute of Bioeconomy Research, 1431 Ås, Norway; Czech University of Life Sciences Prague, Faculty of Forestry and Wood Sciences, EVA 4.0 Unit, Kamýcká 129, CZ – 165 00 Praha 6 – Suchdol, Czech Republic; Entomology Department, Max Planck Institute for Chemical Ecology, 07745 Jena, Germany; Department of Plant Protection Biology, Swedish University of Agricultural Sciences, Alnarp 230 53, Sweden

**Keywords:** Coleoptera, Curculionidae, Scolytinae, bark beetle, long-read sequencing, RNA-Seq, automated annotation, *de novo* sequencing

## Abstract

The Eurasian spruce bark beetle (*Ips typographus* [L.]), is a major killer of spruce forests across the Palearctic. During epidemics, it can destroy over 100 million cubic meters of spruce trees in a single year. Here we report a 236 Mb, highly contiguous *I. typographus* genome assembly using PacBio long-read sequencing. The final phased assembly had a contig N_50_ of 6.65 Mb in 272 contigs and was predicted to contain 23,923 protein-coding genes. Comparative genomic analysis revealed expanded gene families associated with plant cell wall degradation, including pectinases, aspartyl proteases, and glycosyl hydrolases. In today’s forests, increasingly stressed by global warming, this resource can assist in mitigating bark beetle outbreaks by developing novel pest control strategies. Further, this first whole-genome sequence from the genus *Ips* provides timely resources to address important questions about the evolutionary biology and ecology of Curculionidae, the true weevils, one of the largest animal families.

## 1 Introduction

Conifer-feeding bark beetles (Coleoptera; Curculionidae; Scolytinae) are keystone species in forest ecosystems, contributing to wood decomposition and nutrient recycling through direct feeding and the action of beetle-associated microbiota (Edmonds & Eglitis, 1989). However, a few so-called aggressive species can also kill large numbers of healthy trees through pheromone-coordinated mass-attacks once their populations surpass a critical threshold density, quickly transforming entire forest landscapes at tremendous economic and ecological costs (Hlásny et al., 2019; Raffa et al., 2016). Since abiotic factors are important drivers for beetle population growth (Biedermann et al., 2019), outbreaks of aggressive species are expected to increase in frequency and severity due to climate change (Kurz et al., 2008; Marini et al., 2017). Beetle population growth is promoted by increasing temperatures that reduce both winter mortality and developmental time, allowing for the production of additional generations per year. Furthermore, the resistance of conifer trees to beetle attack is compromised during heat and drought stress (Allen et al., 2010).

The Eurasian spruce bark beetle (*Ips typographus* [L.]) is the most serious pest of Norway spruce (*Picea abies* [L.] Karst) and other spruce species in the beetle’s range, currently causing unprecedented forest destruction across the Palearctic region (**Fig. 1**). In Europe, *I. typographus* killed between 2 and 14 million m^3^ spruce annually from 1970 to 2010 (Seidl et al., 2014) but during the hot and dry summer of 2019, it killed more than 110 million m^3^ (Ebner, 2020). Beetle mass-attacks on trees are coordinated by an aggregation pheromone made up of *cis*-verbenol and 2-methyl-3-buten-2-ol. The pheromone is produced by males as they initiate boring in suitable host trees and attracts large numbers of both sexes (Schlyter et al., 1987). The pheromone component *cis*-verbenol is produced from the spruce monoterpene (−)-4*S*-α-pinene (Lindström et al., 1989), whereas 2-methyl-3-buten-2-ol is produced *de novo* by the beetles (Lanne et al., 1989). Pheromone attraction is modulated by other compounds produced by con- and heterospecific beetles as well as by host and non-host trees, highlighting the importance of chemosensation in this bark beetle (Andersson et al., 2010; Schlyter et al., 1989; Zhang & Schlyter, 2004). Due to the beetle’s economic and ecological importance, studies have started to address the molecular biology and function of chemosensation in *I. typographus*, but genetic resources are scarce and have thus far been limited to transcriptomic data (Andersson et al., 2013; Yuvaraj et al., 2020).

**FIGURE 1.**
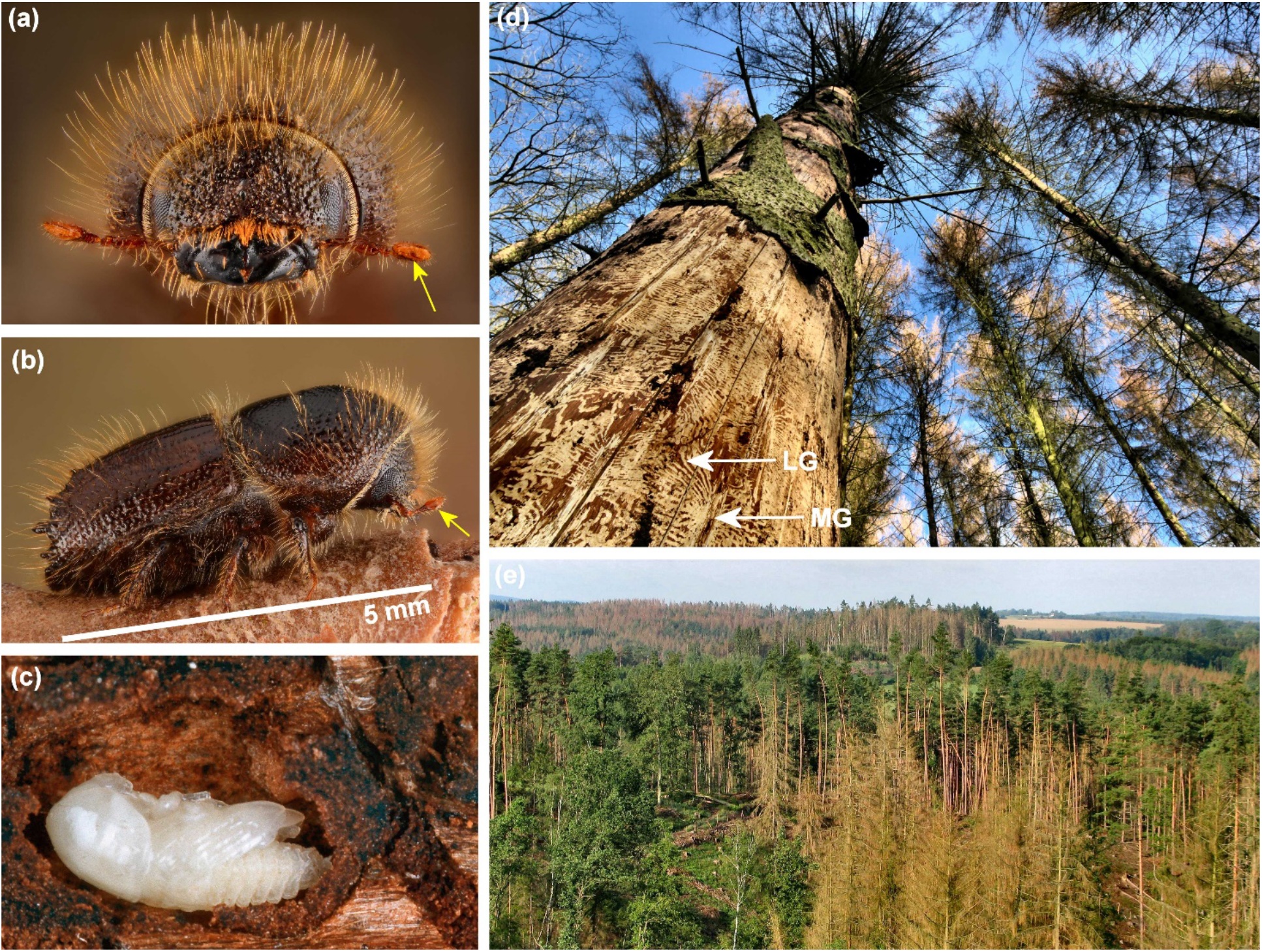
Adult *Ips typographus* front (a) and side view (b) with sensory array on antennae indicated (yellow arrows) and scale bar. (c) Pupae in bark. (d) Damage to Norway spruce (*Picea abies*) at tree level (with two of several hundred galleries indicated with white arrows; MG: conventional mother gallery; LG: winding larval galleries). (e) Damage at landscale level of mixed forest with dead Norway spruce trees, while trees with green crowns are Scots pine or deciduous species. Pictures (a, b, d) from Belgium by Gilles San Martin, (c) by blickwinkel/Alamy Stock Photo, and (e) from Czech Republic by Jan Liška.

Tree-killing bark beetles like *I. typographus* thrive on the inner bark (**Fig. 1a-d**), one of the most nutritionally poor and recalcitrant organic substrates on earth (Filipiak & Weiner, 2017). Abundant carbon-rich bio-polymers present in bark and adjoining sapwood, such as lignin, hemicellulose and cellulose may not be available to bark beetles as nutrients until prior degradation by fungi and other microbes (Kirk & Cowling, 1984). Bark beetles, including *I. typographus*, are associated with a holobiont of diverse microbiota, including gut endosymbionts (Chakraborty et al., 2020a; Chakraborty et al., 2020b), yeasts (Davis, 2015), and ecto-symbionts such as ophiostomatoid fungi that the beetles inoculate into attacked trees (Kirisits, 2007). Once established in the tree, these fungi may serve as food for developing beetles, metabolise conifer chemical defences, and accelerate tree death (Christiansen, 1985; Kandasamy et al., 2019; Zhao et al., 2019).

Although beetles (Coleoptera) constitute the largest order of the Metazoa, only 11 Coleoptera genomes have been published, of which only two belong to members of the 6,000 species strong Scolytinae subfamily (the mountain pine beetle, *Dendroctonus ponderosae* Hopkins, (Keeling et al., 2013) and the coffee berry borer, *Hypothenemus hampei* Ferrari (Vega et al., 2015)). Sequencing of additional Scolytinae genomes would enable comparative genomic studies and shed light on taxon-specific ecological adaptations in this highly specialized and important insect group. The main aim of this study was to assemble and annotate a high-quality genome of *I. typographus*. Secondly, we performed a comparative genomic analysis with 20 other metazoan genomes, specifically investigating *Ips*- and bark beetle-specific gene family expansions. We expect that our high-quality assembly of the *I. typographus* genome will be an important resource for forthcoming fundamental and applied studies on this important insect that endangers entire forest landscapes (**Fig. 1e**).

## 2 Materials and Methods

### 2.1 Insect rearing and nucleic acid extraction

To reduce heterozygosity in the beetles before sequencing, we ran 10 generations of sibling mating. The first mating pairs were taken from a laboratory-reared population of *I. typographus* originating from Lardal, SW Norway. The continuous culture was kept at the Swedish University of Agricultural Sciences in Alnarp, Sweden on local, freshly cut Norway spruce, following the procedures described in Anderbrant et al. (1985). Sexes were separated by pronotum hair density (Schlyter & Cederholm, 1981) and each mating pair had a 28 × 12-14 cm (length × diameter) spruce bolt providing food *ad libitum* (Anderbrant et al., 1985). For each new sibling mating generation, we used siblings from offspring from one or occasionally two bolts (strictly separated), ensuring strict sibling mating. To reduce the risk of breeding failure, the 10 successful sibling mating generations were undertaken by using at least 10 spruce bolts per generation, with little or no observed inbreeding depression effects on fecundity or body mass over time. High molecular weight (HMW) DNA was extracted from around 100 individuals from this 10× inbred population to be used for high-throughput sequencing. Beetles were frozen in liquid nitrogen and pulverised using a pre-chilled mini-mortar and pestle. The resulting powder was transferred with a pre-chilled spatula to a microfuge tube containing lysis buffer and ball bearings. The sample was further homogenised using a Qiagen TissueLyzer by shaking for 1 min at 50 Hz. The optimal method for isolation of good quality HMW DNA from *I. typographus* was using the Qiagen Genomic Tip 100/G kit following the manufacturers’ instructions, except for a prolonged incubation time at 50 °C overnight with gentle agitation using a modified lysis buffer (20 mM EDTA, 100 mM NaCl, 1% Triton^®^ X-100, 500 mM Guanidine-HCl, 10 mM Tris, pH 7.9) replacing the standard Qiagen G2 buffer, and with the addition of Proteinase K to 0.8 mg/ml. This was supplemented with the addition of DNase-free RNase A (20 μg/ml) treatment and incubation for 30 min at 37 °C, followed by centrifugation for 20 min at 12,000× g to pellet insoluble debris. The clarified lysate was then transferred to the QBT buffer-equilibrated Genomic Tip to proceed with the standard protocol.

### 2.2 Genome sequencing, assembly and evaluation

DNA samples were transported to the sequencing facility at the National Genomics Infrastructure SciLife Laboratories in Uppsala, Sweden for library preparation and sequencing on the PacBio RS II platform (Pacific Biosciences). Primary assembly of the PacBio long reads was performed using FALCON-kit v1.3.0 software and haplotype phasing was achieved using FALCON-unzip v1.2.0 with default settings (Chin et al., 2016). Contigs from the final assembly were aligned against each other using MUMmer (Kurtz et al., 2004) and redundant contigs removed. The completeness of the genome was assessed using metrics derived from read alignment and searches of single-copy orthologs. The Benchmarking Universal Single-Copy Orthologs (BUSCO v3.0.2) (Simão et al., 2015) tool was used to search the genome assembly against the insecta_odb9 database of 1,658 genes. In order to obtain an understanding of the chromosomal structure of long contigs, telomeric motifs were identified using the script FindTelomeres.py (https://github.com/JanaSperschneider/FindTelomeres). Genome size estimation was performed via *k*-mer counting with Jellyfish v2.3.0 (Marçais & Kingsford, 2011) using GenomeScope 2.0 (Ranallo-Benavidez et al., 2020) from a genome survey of 364 million 150 bp Illumina reads equivalent to approximately 230-fold coverage.

### 2.3 Genome annotation

A custom repeat library was produced from the genome assembly using RepeatModeler v1.0.11 (http://www.repeatmasker.org/RepeatModeler/). This custom library, together with the Repbase library, was used with RepeatMasker v4.0.8 (http://www.repeatmasker.org/) to soft mask the assembly prior to annotation. RNA-Seq data from multiple beetle life stages and tissues (**Supplementary Table 1**) were aligned to the genome assembly using HiSat2 v2.1.0 (Kim et al., 2015) and annotated using StringTie v1.3.3 (Pertea et al., 2015). The alignment features were passed to BRAKER2 (Hoff et al., 2015) as hints for training an AUGUSTUS model using GeneMark-ET. The transcriptome assembly from the above RNA-Seq data was used together with the published protein sequences from the genomes of three species of Coleoptera namely, *D. ponderosae* (Scolytinae), *Anoplophora glabripennis* Motschulsky (Cerambycidae), *Tribolium castaneum* Herbst (Tenebrionidae), and also the protein sequences from a further eight high-quality genomes, including *Drosophila melanogaster* Meigen (Diptera) and *Homo sapiens* Linné, as evidence for homology-based gene prediction using MAKER3 (Holt & Yandell, 2011), as outlined in **Supplementary Fig. 1**. Because transcriptome data is prone to misassembly and chimaeras it was only used for evidence and not for prediction. Protein sequences identified from the BUSCO searches were passed to the MAKER3 pipeline for training SNAP (Korf, 2004) along with the BRAKER2 trained AUGUSTUS model for *ab initio* predictions, using the HiSat2 RNA-Seq alignments as transcript-based evidence. Gene model predictions were retained if they were greater than 33 amino acids in length and were supported by at least one form of supporting evidence of either (i) a BLASTp match (E-value < 10^-10^) to the non-redundant protein GenBank database, (ii) coverage by RNA-Seq data alignment, or (iii) containing a hit to the Pfam database. Several iterations of MAKER3 were performed to improve upon the prediction in subsequent runs.

### 2.4 Comparative genomics and phylogenomics

In order to identify expansions of gene families within the *I. typographus* genome, we searched the protein-coding sequences for Pfam (Finn et al., 2014) domains to assign gene function. We used HMMER v3.1 (Potter et al., 2018) (hmmscan) to search the Pfam A database (release 32.0) for 13,841 different domains of 20 different species of Metazoa. Counts of each domain were collated for each species and domains that occurred multiple times in a protein sequence were counted only once. A Fisher’s exact test was then conducted iteratively using R, comparing the number of counts in Pfam families found in an individual genome, normalised by the total gene count for that species, against the background, which we defined as the average of the counts in the remaining species. Multiple testing corrections were done using the FDR method in R for all calculated *p*-values. A Pfam domain was considered expanded in *I. typographus* if it showed a corrected *p*-value < 0.05.

Protein sequences from *I. typographus, D. ponderosae, T. castaneum, H. hampei* and *A. glabripennis* were compared for orthology using all-against-all alignments (*£*-value of 10^-5^) and clustered using OrthoVenn2 (Xu et al., 2019) with an inflation value of 1.5. Phylogenetic inference from orthologous genes was undertaken by comparing the *I. typographus* gene set with the proteincoding sequences from 11 other species of Coleoptera (**Supplementary Table 2**) using OrthoFinder v2.4.0 (Emms & Kelly, 2019). Orthologous gene groups were used to build a rooted species tree that was visualised using FigTree v1.4.4 https://github.com/rambaut/figtree/releases. Presence of genes associated with detoxification and pesticide resistance in sequenced species of Coleoptera were identified based on Pfam domains contained within their respective protein sequences.

### 2.5 Transcriptome assembly and quantification of gene expression

Transcriptome libraries from different *I. typographus* life stages and tissue types (larval stages 1-3, pupae, adult beetle, male head, female head, callow male gut and fat body, callow female gut and fat body) were prepared at the Czech University of Life Sciences, Prague (**Supplementary Table 1**). Whole larvae, pupae and adult *I. typographus* were sampled along with discrete tissues that were dissected from adults and pooled under sterile conditions for RNA extraction. Total RNA was purified using the PureLink™ RNA Kit from Ambion (Invitrogen) following the manufacturer’s protocol. The integrity of the purified total RNA was checked using agarose gel electrophoresis and the 2100 Bioanalyzer system (Agilent Technologies, Inc). Total RNA samples with an integrity number (RIN) > 7 were selected for sequencing. Total RNA was quantified using a Qubit 2.0 Fluorometer (Thermo Fisher Scientific) with an RNA HS Assay Kit (Invitrogen) before sending for sequencing at Novogene, China, using the Illumina platform to generate a minimum of 30 million 150 bp paired-end reads per sample. Five biological replicates for each sample type were sequenced. Raw reads were processed using Trimmomatic v (Bolger et al., 2014) and assembled using Trinity v2.8.2 (Haas et al., 2013) with the default parameters except for --normalize_max_read_cov 100 -- min_kmer_cov 3 --min_glue 3 --KMER_SIZE 23. Expression levels were measured by aligning quality-processed RNA-Seq reads to the genome assembly using HiSat2 v2.1.0. Aligned reads were sorted with SAMtools v1.5 (Li et al., 2009) and counts reported as transcripts per one million mapped reads (TPM) using StringTie v1.3.3. TPM values visualised in heatmaps were transformed to log2 (TPM+1) and normalised across tissues using the scale function in R.

## 3 Results and Discussion

### 3.1 Genome sequencing, assembly and completeness assessment

A highly inbred line of *I. typographus* was successfully produced via full-sib mating for 10 generations. Whole-genome sequencing yielded over 4.5 million subreads totalling more than 50 gigabases of sequence equivalent to 211-fold coverage. The subread N_50_ was 19.3 kb with a mean read length of 12.4 kb. Estimation of genome size using a kmer-counting approach suggested a total haploid size of 221.9 Mb (**Fig. 2**), comparable with the mountain pine beetle (*D. ponderosae*) genome assembly of 204 Mb (Keeling et al., 2013). The final, phased genome sequence was 236 Mb in total length and comprised 272 contigs with an N_50_ of 6.65 Mb; the longest contig being 16.9 Mb in size (**Table 1**). Seventy-eight percent of the genome assembly was contained in 36 contigs that were greater than 1 Mb in size (**Supplementary Table 3**). The five largest contigs were all greater than 10 Mb in length.

**FIGURE 2.**
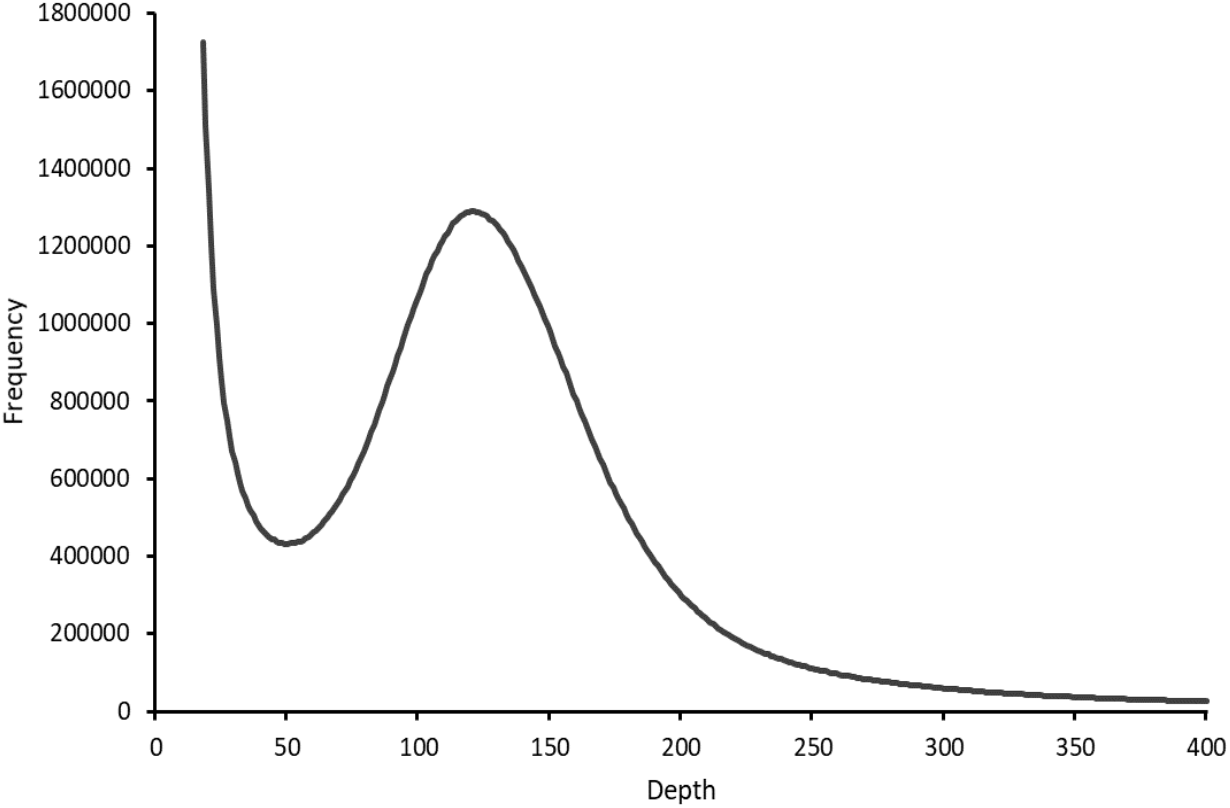
*K*-mer distribution (*k* = 19) of Illumina sequencing reads of *Ips typographus*. The peak *k*-mer depth was 124, giving an estimated haploid genome length of 221,929,798 bp. The presence of a single peak suggests a highly homozygous genome.

**TABLE 1.** Assembly statistics of the *Ips typographus* genome and comparisons with two other species of Coleoptera, *Dendroctonus ponderosae* and *Tribolium castaneum*.

| Statistic | ltyp1.0 (this study) | DendPond_male_1.0<sup>1</sup> | Tcas5.2<sup>2</sup> |
| --- | --- | --- | --- |
| Genome size (Mb) | 236.82 | 252.85 | 165.93 |
| Number of contigs | 272 | 8188 | 2081 |
| Contig N50 (Mb) | 6.65 | 0.63 | 15.27 |
| Contig L50 | 12 | 87 | 5 |
| Max. contig length (Mb) | 16.86 | 4.16 | 31.38 |
| Min. contig length (kb) | 20.29 | 1.00 | 0.25 |
| GC (%) | 35.21 | 35.91 | 33.86 |
| BUSCO Completeness (insecta_odb9) | 98.6% | 95.4% | 99.3% |
| Complete and single-copy | 1535 (92.6%) | 1489 (89.8%) | 1643 (99.1%) |
| Complete and duplicated | 99 (6%) | 93 (5.6%) | 4 (0.2%) |
| Fragmented | 15 (0.9%) | 49 (3.0%) | 8 (0.5%) |
| Missing | 9 (0.5%) | 27 (1.6%) | 3 (0.2%) |
| RNA-Seq alignment rate (% total) | 93.56 | - | - |
| RNA-Seq alignment rate (% concordantly exactly 1 time) | 83.73 | - | - |

Note: Comparison genomes downloaded from GenBank; <sup>1</sup>*Dendroctonus ponderosae* GCA\_000355655.1; <sup>2</sup>*Tribolium castaneum* GCA\_000002335.3.

Telomeric motifs are regions of repetitive sequences of DNA that indicate the ends of chromosomes. These motifs were identified at the ends of eight different contigs (five forward strands, three reverse strands). Teleomeric motifs were located at one end of the five largest contigs (**Supplementary Table 4**) suggesting that each of these largest contigs start or finish at the end of a chromosome. The overall GC content was 33.21%. Approximately 28.2% of the genome contained repetitive sequence when masked using a custom library. This was slightly higher than in *D. ponderosae* (between 17% and 23%) (Keeling et al., 2013), though lower than in *T. castaneum* (42%) (Richards et al., 2008; Wang et al., 2008). Analysis of completeness of the draft genome using the BUSCO tool revealed that 1,649 (99.5%) of the 1,658 genes in the insecta_odb9 database could be identified as present either partial or complete. Only nine genes (0.5%) were considered missing from the assembly. The *I. typographus* assembly statistics were of a quality comparable with the release of the enhanced *T. castaneum* genome sequence (Herndon et al., 2020) and were considerably higher than most other Coleoptera genomes published to date. Thus, our evaluation indicates that the *de novo* assembly of the *I. typographus* genome is of high quality. Comparisons of contig sizes with other species and the presence of multiple telomeres suggest some of the largest contigs contained in this assembly may be approaching chromosome scale.

### 3.2 Genome annotation

Our approach to annotate the *I. typographus* genome exploited extensive transcriptomic resources and numerous publicly available protein sequences using *ab initio* and homology-based predictors. The final set of models consisted of 23,923 protein-coding genes (**Table 2**) with the majority (> 77%) assigned an annotation edit distance (AED) score of 0.5 or less (**Supplementary Fig. 3**). At least 84% (20,094) of the gene models were supported by alignments to our RNA-Seq read data. Searches of the annotated predicted protein sequences using BUSCOs from the insecta_odb9 libraries identified 1,584 (95.5%) of the 1,658 protein sequences in the Insecta dataset, with only 38 (2.3%) sequences missing (**Supplementary Fig. 4**). Comparing the outcome of BUSCO searches with the published Scolytinae (*D. ponderosae* and *H. hampei*) gene models, the *I. typographus* predictions had similar levels of completeness as these and were also comparable with the higher-quality coleopteran genome assemblies, such as the two other wood-feeding beetles (*Agrilus planipennis* and *Anoplophora glabripennis;* **Supplementary Fig. 4**). Moreover, read alignment of RNA-Seq data to annotated regions of the draft genome resulted in an average of 83.7% of reads mapping concordantly in pairs to the gene space. An overall average of 93.6% mapped reads indicates that the majority of transcripts captured in our RNA-Seq data can be found and are intact in the *I. typographus* gene set. Of the 23,923 predicted protein sequences, 59% contained Pfam domains. The number of gene models for *I. typographus* was considerably higher than that reported for *D. ponderosae* (13,088) (Keeling et al., 2013), though comparable with the annotation of *A. glabripennis* (McKenna et al., 2016) and *H. hampei* (Vega et al., 2015) with 22,035 and 19,222 predicted protein-coding genes, respectively. Taken together, these statistics suggest the production of a comprehensive set of gene models for *I. typographus*.

**TABLE 2.**
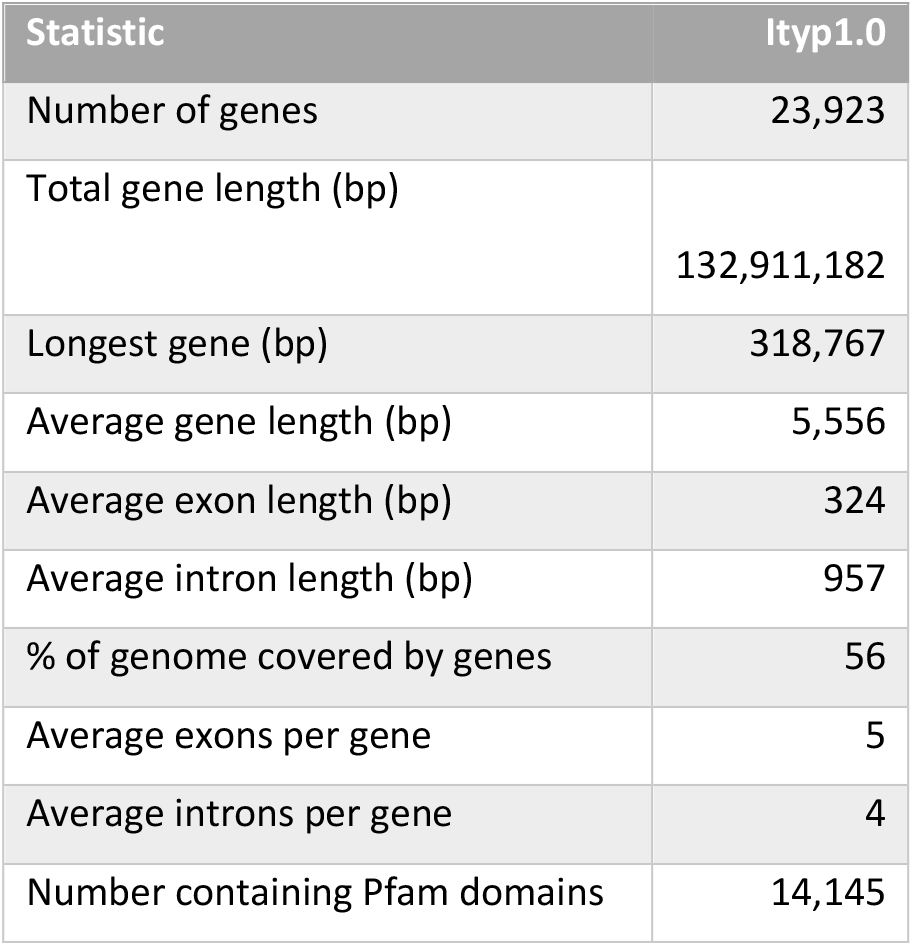
Statistics of the predicted gene models from the *Ips typographus* draft assembly (Ityp1.0).

### 3.3 Orthology and comparative analysis with other Coleopteran genomes

Comparison of orthologous genes shared among three beetle species closely related to *I. typographus* showed a core set of 6,991 shared gene clusters and a unique set of 811 clusters that were specific to *I. typographus* (**Fig. 3a**). Gene ontology enrichment analysis of these *Ips*-specific gene clusters revealed nine enriched categories, including DNA and RNA binding, regulation, and signalling (**Supplementary Table 5**). A total of 22 gene families were significantly expanded in *I. typographus* when compared with the 11 other published coleopteran genomes and a selection of other model invertebrates. A selection (17) of these expanded gene families are shown in **Fig. 3b**. The limited number of significantly expanded gene families discovered in *I. typographus* in this analysis reflects the comprehensive array of beetle species included in the dataset. Using this comprehensive approach, we were able to highlight gene families that appear distinctly relevant to the ecology of *I. typographus* and/or conifer-feeding bark beetles. A phylogenetic tree inferred from orthologous gene families shows the phylogenetic position of *I. typographus* relative to the other coleopterans included in this study (**Fig. 3c**).

**FIGURE 3.**
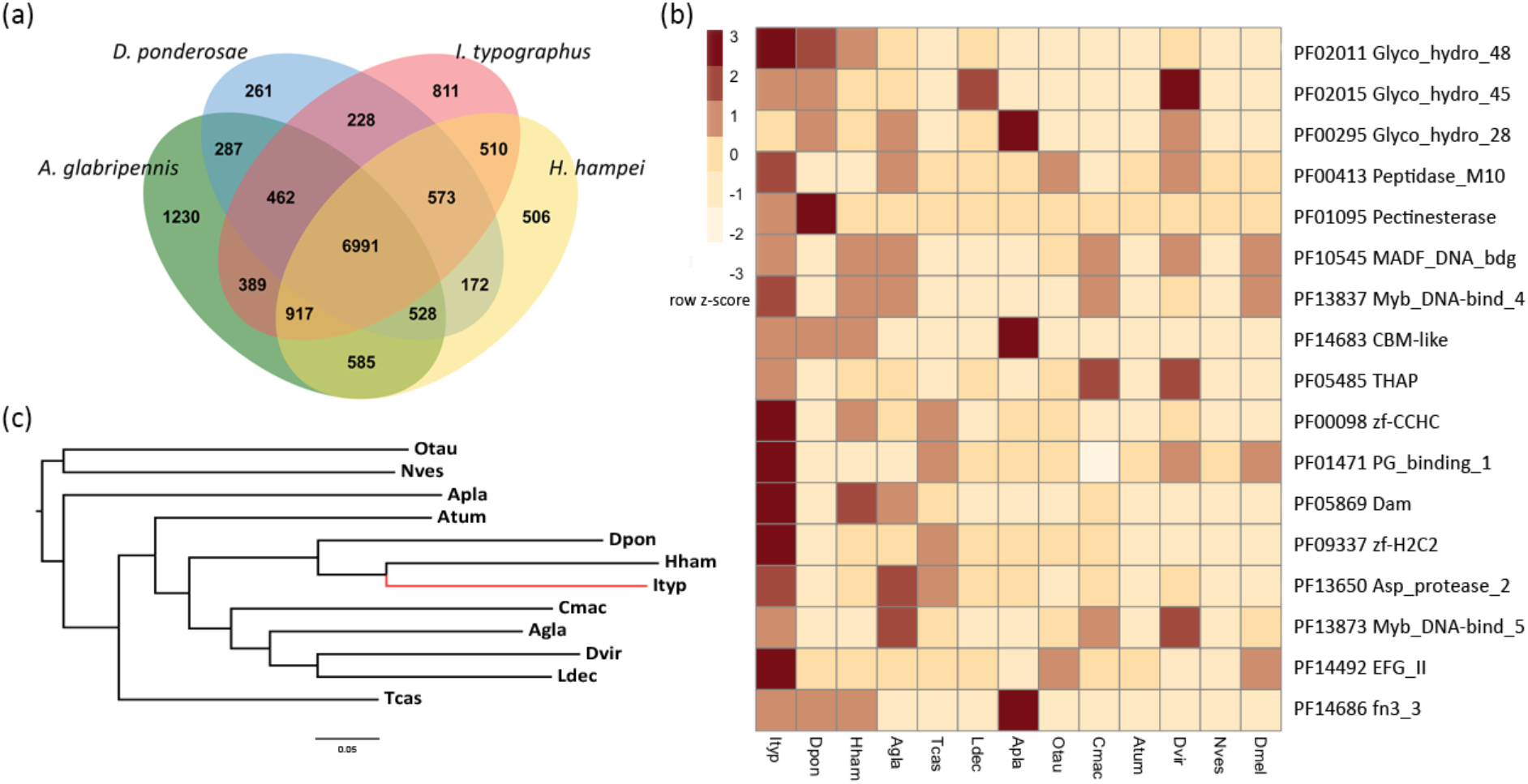
(a) Venn diagram of orthologous genes shared between *I. typographus, D. ponderosae, H. hampei and A. glabripennis*. (b) Expanded protein families in *I. typographus* compared with other sequenced Coleoptera species and *D. melanogaster*. Cell colours indicate the number of standard deviations from the mean of each domain count for all species. (c) Phylogenetic relationship of 11 Coleoptera species inferred from orthologous gene groups. Atum, *Aethina tumida*; Apla, *Agrilus planipennis;* Agla, *Anoplophora glabripennis;* Cmac, *Callosobruchus maculatus;* Dpon, *Dendroctonus ponderosae;* Dvir, *Diabrotica virgifera;* Hham, *Hypothenemus hampei;* Ityp, *Ips typographus;* Ldec, *Leptinotarsa decemlineata;* Nves, *Nicrophorus vespilloides;* Otau, *Onthophagus taurus;* Tcas, *Tribolium castaneum*; Dmel, *Drosophila melanogaster*.

Feeding on the stem bark of trees necessitates the ability to metabolize plant cell walls using endogenous enzymes or relying on the actions of associated microbiota. Notably, genes containing domains associated with plant cell wall degrading enzymes (PCWDEs) were significantly expanded in the *I. typographus* genome. Large expansions of PCWDEs were previously reported in the bark beetle *D. ponderosae* and the polyphagous cerambycid wood-borer *A. glabripennis* (Keeling et al., 2013; McKenna et al., 2016). In particular, three families of glycosyl hydrolase (GH) enzymes were expanded in the conifer-feeding bark beetles *I. typographus* and *D. ponderosae* (**Fig. 3b**). Genes containing the Pfam domain GH48 (PF02011) were most abundant in these species, possessing eight (*I. typographus*) and six (*D. ponderosae*) proteins containing this domain. GH48 enzymes are reducing end acting cellobiohydrolases and have been identified in several polyphagous insect species, but chiefly among the two Coleopteran superfamilies Chrysomeloidea and Curculionoidea (Eyun et al., 2014). A detailed phylogenetic analysis of the Coleoptera and their GH gene repertoire has been reported previously (McKenna et al., 2019). The most abundant expression of these genes in *I. typographus* occurred predominantly in the fat body, with some genes being specifically expressed during different larval stages, but none in pupae (**Fig. 4a**). This finding is surprising since these genes typically are highly expressed in guts, and further experiments are needed to investigate functional roles of these proteins in the fat body. GH enzymes are also commonly found in bacteria and more rarely in fungi (Berger et al., 2007) and their presence in insects is thought to be the result of horizontal gene transfer events from bacteria (Chu et al., 2015; Eyun et al., 2014; McKenna et al., 2019; Pauchet & Heckel, 2013). Fungal symbionts assist *I. typographus* in colonising spruce trees and may play essential roles in beetle nutrition and weakening host tree defences (Kandasamy et al., 2019; Zhao et al., 2019). GH48 enzymes are thought to assist in the degradation of fungal chitin (Fujita et al., 2006), which may be important for *I. typographus* as laboratory bioassays have shown a strong feeding preference of immature adults for media colonized by symbiotic fungi (Kandasamy et al., 2019). Three separate tandem duplication events appear to have occurred in *I. typographus* since the establishment of GH48 genes in the Curculionoidea, seen as three pairs of genes occurring adjacent to each other in the genome (sequentially numbered gene models are found adjacently in the genome). Phylogenetic analysis of all the GH48 domain-containing genes in the 11 species of Coleoptera analysed in this study highlights that the eight genes from *I. typographus* share the highest similarity with those from the other Scolytinae species *D. ponderosae* and *H. hampei* (**Fig. 4b**).

**FIGURE 4.**
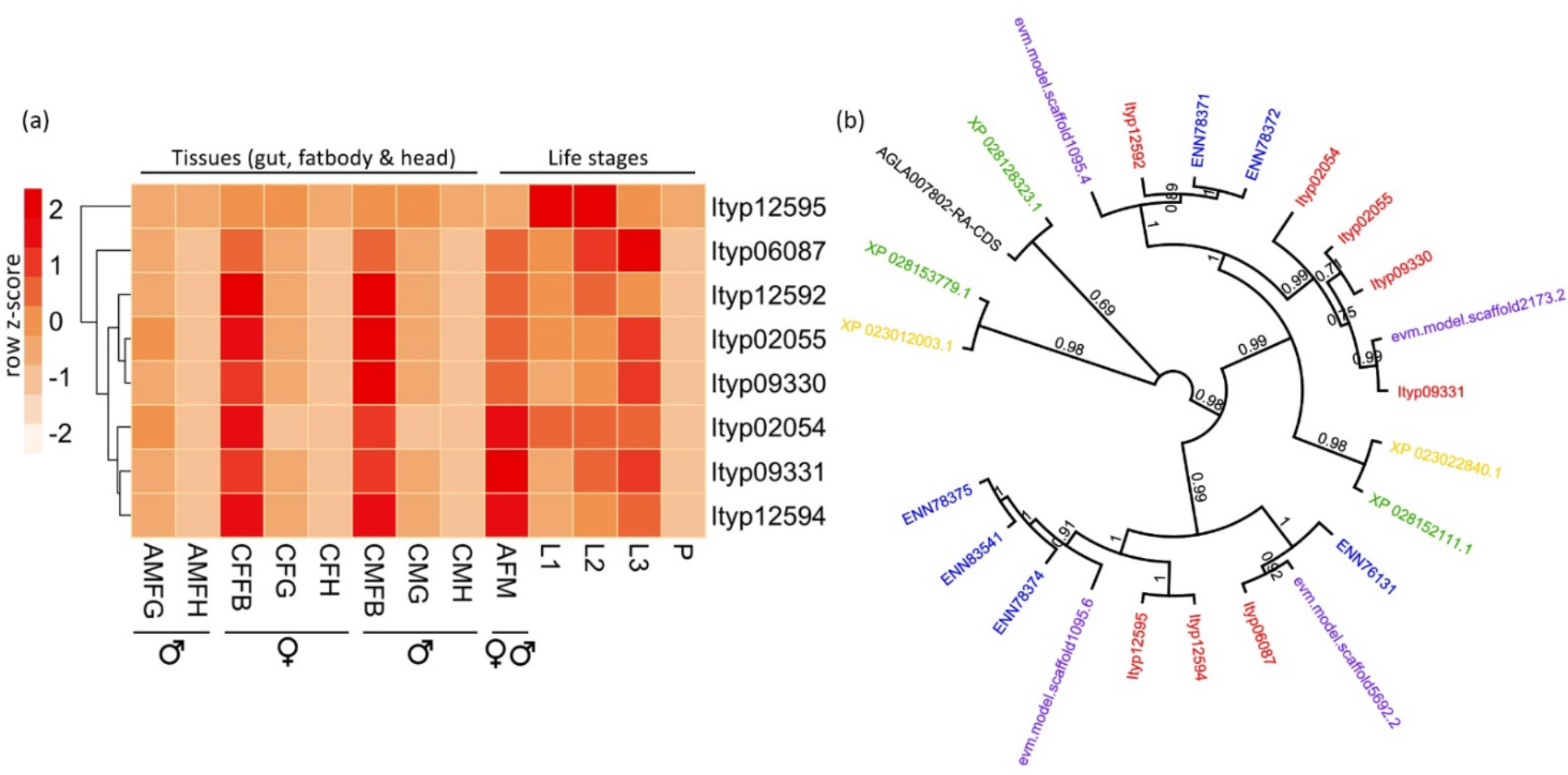
(a) Expression of the expanded GH48 domain (PF02011) containing proteins in 12 *I. typographus* transcriptomes. AMFG, fed adult male gut; AMFH, fed adult male head; CFFB, callow female beetle fat body; CFG, callow female beetle gut; CFH, callow female beetle head; CMFB, callow male beetle fat body; CMG, callow male beetle gut; CMH, callow male beetle head; AFM, adult beetle (male & female); L1, larvae stage 1; L2, larvae stage 2; L3, larvae stage 3; P, Pupae. Cell colours indicate the number of standard deviations from the mean expression level. (b) Phylogenetic tree of GH48 domain-containing proteins in *Ips typographus* (red) and five other Coleoptera species (blue, *Dendroctonus ponderosae*; purple, *Hypothenemus hampei*; black, *Anoplophora glabripennis*; green, *Diabrotica virgifera*; yellow, *Leptinotarsa decemlineata*).

The *I. typographus* genome contains 33 aspartyl protease (PF13650) domaincontaining genes. Although these genes are also expanded in the wood-borer *A. glabripennis*, this is more than any of the other species included in this study, and clearly distinguishes *I. typographus* from *D. ponderosae* that has only a single gene containing this domain. Aspartyl protease is a digestive protease and has been implicated in proteolytic digestion in insects (Santamaria et al., 2015). Both pectinesterase (PF01095) and polysaccharide lyase (PF14683) gene families were found enriched in the three bark beetles. GH28 genes were also expanded in the bark beetles, but also in the wood borers *A. glabripennis* and especially the emerald ash borer, *Agrilus planipennis* (Buprestidae). These gene families are known to be involved in the degradation of pectin, which is a major component of primary plant cell walls. In several beetle species (*D. ponderosae, H. hampei, A. glabripennis*, and *L. decemlineata*) there is evidence suggesting a fungal origin of the GH28 genes (Schoville et al., 2018, and references therein). Expansions of genes involved in cell wall degradation may extend the capacity of these species to digest their recalcitrant host plants efficiently.

Searches of protein families within 12 species of Coleoptera revealed differences in the repertoire of genes important for detoxification and pesticide resistance. Interestingly, comparison between *I. typographus* and *D. ponderosae* revealed many similarities in gene family abundance (**Table 3**). However, these two conifer-infesting bark beetles did not display an enrichment for gene families involved in xenobiotic metabolism when compared to other agriculturally important Coleopterans. In particular, the number of cytochrome P450 genes in the *I. typographus* genome (86 genes) is lower than in all the other species included in our analysis, except for *H. hampei* (67 genes). However, the number of P450 genes in *D. ponderosae* is only marginally higher (93 genes), and many of these genes were shown to form species-specific expansions mainly within the CYP6 and CYP9 subfamilies (Keeling et al., 2013). Hence, it is possible that Scolytinae specialists on conifers detoxify host defences (such as abundant conifer monoterpenes and phenolics) using a smaller, but specialized, repertoire of P450 enzymes, as compared to generalist beetles feeding on diverse angiosperms. Conifers in the pine family (Pinaceae) are rich in preformed terpenes and conifer specialists might therefore be expected to have numerous P450s to detoxify these. However, *I. typographus* seems to be less tolerant to terpenes than species of *Dendroctonus* (Everaerts et al., 1988). This may appear paradoxical considering its lifestyle, but *I. typographus* may have a strategy of quickly reducing the levels of tree defences by mass-attack and assistance from fungal symbionts (Franceschi et al., 2005). Correspondingly, Norway spruce trees have less preformed defences and rely more on capability for induced response for their resistance against bark beetle attacks (Schiebe et al., 2012).

**TABLE 3.**
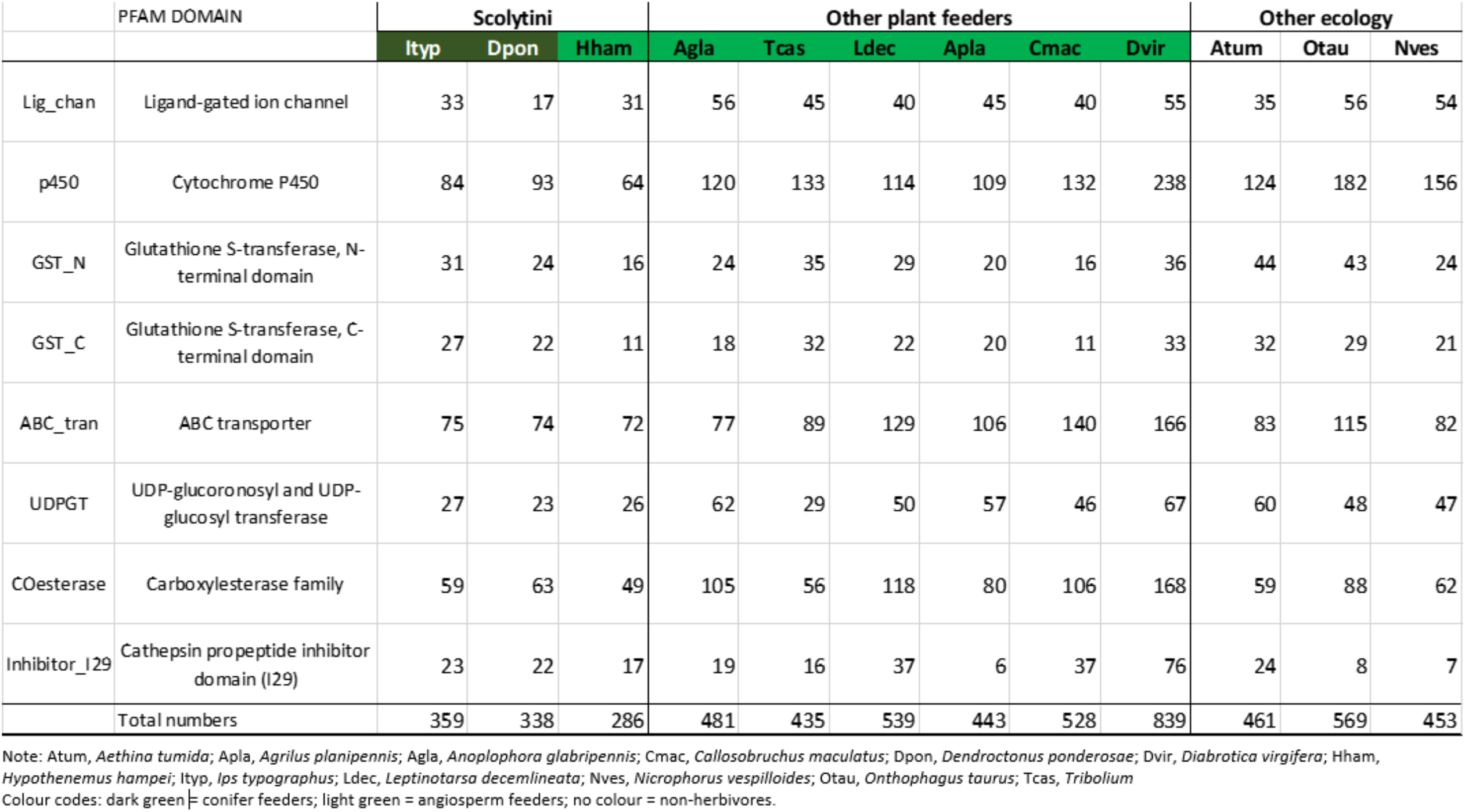
Comparison of protein families associated with detoxification and pesticide resistance between *Ips typographic* and 11 other Coleoptera species. The number of proteins containing the specific Pfam domain for each family is given.

## 4 Conclusion

We assembled and annotated the genome of the Eurasian spruce bark beetle *Ips typographus*, an important pest of spruce across the Palearctic. This highly contiguous, phased assembly comprises 236 Mb of sequence in 272 contigs with an N_50_ of 6.65 Mb and contains 23,923 annotated genes. Our analyses of this assembly suggest gene family expansions of primarily plant cell wall degrading enzymes, of which aspartyl protease domain-containing genes stand out as particularly expanded. P450 enzyme-encoding genes involved in hormone synthesis and catabolism of toxins are reduced in numbers. This first whole-genome sequence from the genus *Ips* provides timely resources for studies addressing important questions about the evolutionary biology and ecology of the true weevils, for population genetic studies, and the management of a forest pest of increasing importance due to heat and drought stress of trees in the Anthropocene.

## ACKNOWLEDGEMENTS

Dr Jan Bily (Czech University of Life Sciences, Prague) is acknowledged for technical support during the preparation of RNA samples for tissue transcriptome study. Infrastructural support and salary for A.R., E.G-W., and F.S. was obtained from project EXTEMIT-KCZ.02.1.01/0.0/0.0/15_003/0000433 financed by OP RDE at Czech University of Life Sciences, Prague. M.N.A. was funded by the Swedish Research Council FORMAS (grants #217-2014-689 and #2018-01444). C.L. acknowledges support from the Swedish Research Council VR (#2017-03804). P.K. was funded by the Research Council of Norway (grant #249958/F20).

## AUTHOR CONTRIBUTIONS

D.P. performed the genome annotation, all bioinformatic analyses and wrote the manuscript, with M.N.A contributing to the initial draft. M.N.A. and F.S. conceived, initiated, and designed the study with E.G-W. and P.K. F.S. performed the sib-mating process with A.C. H.V. did DNA extraction. C.L. provided resources for the bioinformatics. A.R. and E.G-W. did tissue transcriptome sequencing for genome annotation. All authors commented on, improved and approved the final manuscript.

## DATA AVAILABILITY STATEMENT

This Whole Genome Shotgun project has been deposited at DDBJ/ENA/GenBank. All sequence data relating to this study will become available under the BioProject accession number PRJNA671615. For the BioRxiv-submitted pre-publication version of this paper the data are available by directly contacting the corresponding author (M.N.A).

