## Supplementary Material for "A highly contiguous genome assembly of a major forest pest, the Eurasian spruce bark beetle *Ips typographus*"

### Supporting Information

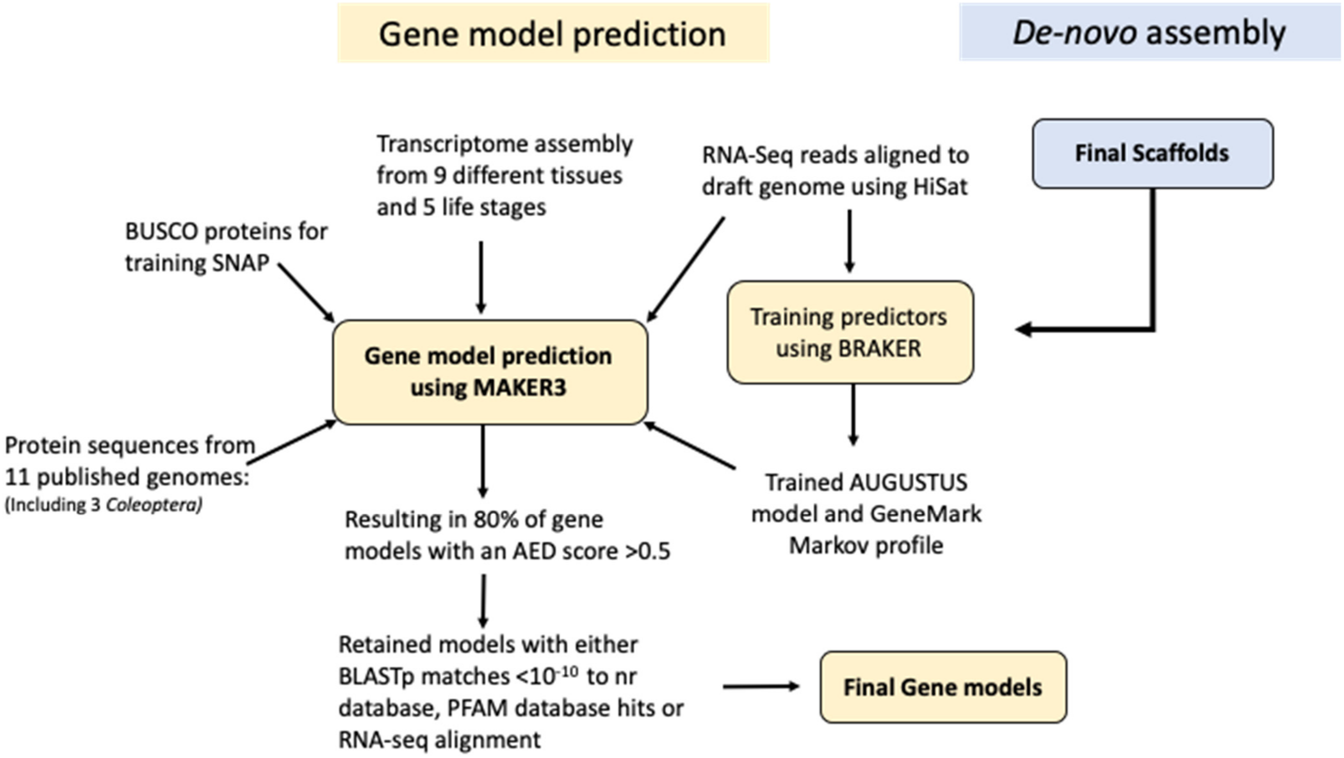


**Supplementary Figure 1**. Workflow diagram of the gene model prediction pipeline for the *I. typographus* genome.

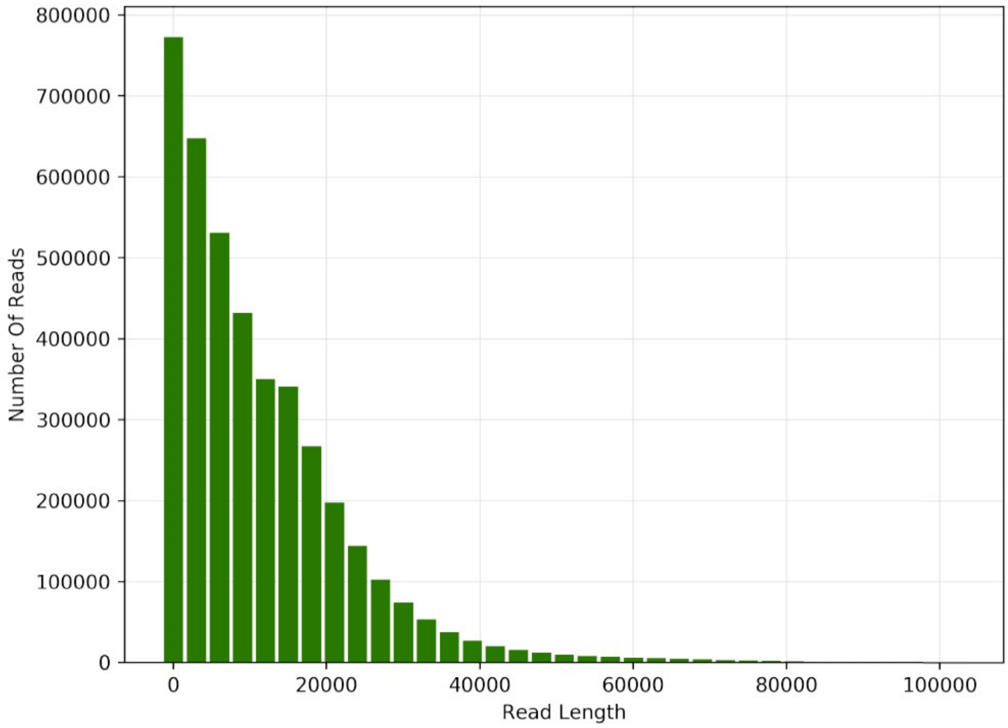


**Supplementary Figure 2.** Read length distribution from PacBio sequencing of the *I. typographus* genome.

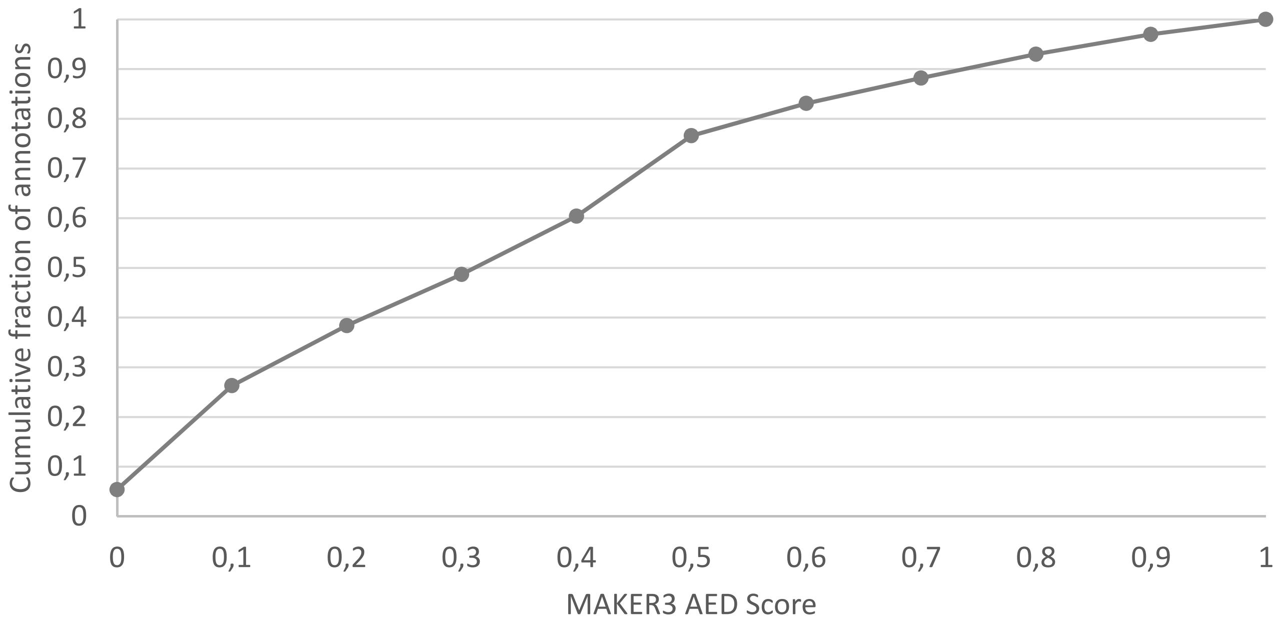


**Supplementary Figure 3.** Cumulative Annotation Edit Distance (AED) scores for gene models produced using the MAKER3 pipeline. Almost 80% of the gene models have an AED score of 0.5 or less. The AED score is a metric assigned to a gene model by MAKER3 and is a measure of the degree of fit that model has with the supporting evidence.

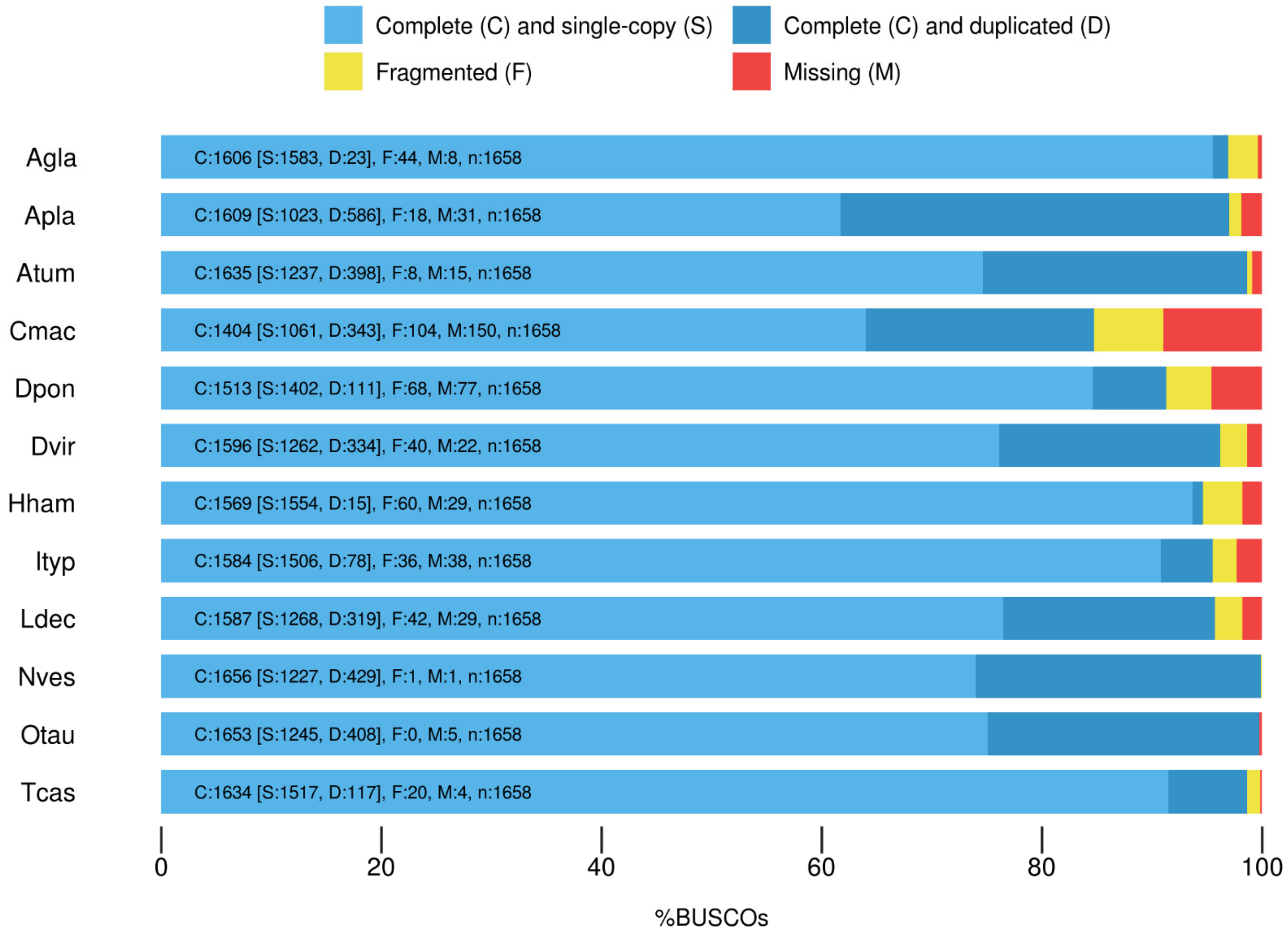


**Supplementary Figure 4.** Completeness estimates of *I. typographus* (Ityp) gene models and comparison with other published coleopteran gene sets using BUSCO tool searches against the Insecta dataset of 1,658 genes. Atum, *Aethina tumida*; Apla, *Agrilus planipennis*; Agla, *Anoplophora glabripennis*; Cmac, *Callosobruchus maculatus*; Dpon, *Dendroctonus ponderosae*; Dvir, *Diabrotica virgifera*; Hham, *Hypothenemus hampei*; Ityp, *Ips typographus*; Ldec, *Leptinotarsa decemlineata*; Nves, *Nicrophorus vespilloides*; Otau, *Onthophagus taurus*; Tcas, *Tribolium castaneum.*

**Supplementary Table 1.** Details of RNA-Seq samples used in this study.

| Sample ID | Description | Number of PE reads | % Mapping to genome |
|---|---|---|---|
| L1 | larvae stage 1 | 171,286,469 | 95.76% |
| L2 | larvae stage 2 | 159,867,283 | 94.62% |
| L3 | larvae stage 3 | 159,565,350 | 93.79% |
| P | pupae | 160,837,226 | 93.98% |
| AFM | adult beetle (male & female) | 181,374,351 | 93.59% |
| AMFG | fed adult male gut | 187,490,572 | 93.25% |
| AMFH | fed adult male head | 173,389,571 | 92.18% |
| CFFB | callow female beetle fat body | 245,181,256 | 93.95% |
| CFG | callow female beetle gut | 189,231,418 | 95.26% |
| CFH | callow female beetle head | 159,092,068 | 89.57% |
| CMFB | callow male beetle fat body | 184,072,277 | 93.12% |
| CMG | callow male beetle gut | 174,127,755 | 94.95% |
| CMH | callow male beetle head | 188,654,217 | 92.25% |

**Supplementary Table 2.** Summary of published genomes used in this study.

| Coleoptera > Polyphaga | | | Taxonomy | | | Genome Stats | | |
|---|---|---|---|---|---|---|---|---|
| **Species** | **ID** | **# Prot seqs** | **Superfamily** | **Subfamily** | **Common Name** | **Scaffold N50** | **Total Size** | **Genbank Accession** |
| *Aethina tumida* | *Atum* | 17,463 | Cucujoidea | Nitidulinae | Small hive beetle | 298,879 | 234 Mb | GCF_001937115.1 |
| *Agrilus planipennis* | *Apla* | 22,159 | Buprestoidea | Agrilinae | Emerald ash borer | 1,113,421 | 353 Mb | GCF_000699045.2 |
| *Anoplophora glabripennis* | *Agla* | 22,343 | Chrysomelidae | Lamiinae | Asian longhorned beetle | 678,234 | 706 Mb | GCF_000390285.2 |
| *Callosobruchus maculatus* | *Cmac* | 31,345 | Chrysomelidae | Bruchinae | Cowpea weevil | 212,245 | 1.007 Gb | GCA_900659725.1 |
| *Dendroctonus ponderosae* | *Dpon* | 13,457 | Curculionoidea | Scolytinae | Mountain pine beetle | 628,732 | 252 Mb | GCF_000355655.1 |
| *Diabrotica virgifera virgifera* | *Dvir* | 28,061 | Chrysomeloidea | Galerucinae | Western corn rootworm | 489,108 | 2.418 Gb | GCF_003013835.1 |
| *Hypothenemus hampei* | *Hham* | 19,222 | Curculionoidea | Scolytinae | Coffee berry borer | 44,715 | 162 Mb | GCA_013372445.1 |
| ***Ips typographus*** | ***Ityp*** | **23,937** | **Curculionoidea** | **Scolytinae** | **Eurasian spruce bark beetle** | **6,654,004** | **236 Mb** | **This study** |
| *Leptinotarsa decemlineata* | *Ldec* | 19,038 | Chrysomeloidea | Chrysomelinae | Colorado potato beetle | 139,046 | 641 Mb | GCF_000500325.1 |
| *Nicrophorus vespilloides* | *Nves* | 19,577 | Staphylinoidea | Nicrophorinae | Burying beetle | 122,407 | 195 Mb | GCF_001412225.1 |
| *Onthophagus taurus* | *Otau* | 21,668 | Scarabaeoidea | Scarabaeinae | Taurus scarab | 337,157 | 267 Mb | GCF_000648695.1 |
| *Tribolium castaneum* | *Tcas* | 18,534 | Tenebrionoidea | | Red flour beetle | 4,456,720 | 165 Mb | GCF_000002335.3 |

**Supplementary Table 3.** Contig size statistics from PacBio sequencing of the *I. typographus* genome.

| Minimum contig length | Number of contigs | Total contig length |
|---|---|---|
| All | 272 | 236,816,287 |
| 10 KB | 272 | 236,816,287 |
| 25 KB | 252 | 236,352,854 |
| 50 KB | 190 | 234,082,415 |
| 100 KB | 164 | 232,251,227 |
| 250 KB | 101 | 222,160,924 |
| 500 KB | 72 | 212,556,464 |
| 1 MB | 36 | 186,489,602 |
| 2.5 MB | 20 | 161,077,642 |
| 5 MB | 14 | 136,853,431 |
| 10 MB | 5 | 71,627,877 |

**Supplementary Table 4.** Identification of telomeric regions ($TTAGGG_N$) in the *I. typographus* genome assembly.

| Contig ID | Location | Contig Size (Mb) |
|---|---|---|
| IpsContig1 | Reverse (end of sequence) | 16.86 |
| IpsContig2 | Forward (start of sequence) | 16.77 |
| IpsContig3 | Forward (start of sequence) | 14.31 |
| IpsContig4 | Forward (start of sequence) | 12.83 |
| IpsContig5 | Reverse (end of sequence) | 10.84 |
| IpsContig23 | Forward (start of sequence) | 2.1 |
| IpsContig51 | Forward (start of sequence) | 0.76 |
| IpsContig93 | Reverse (end of sequence) | 0.28 |

**Telomeres found: 8 (5 forward, 3 reverse)**

**Supplementary Table 5.** Gene ontology enrichment of the 811 orthologous gene clusters unique to *Ips typographus* when compared with *Dendroctonus ponderosae*, *Hypothenemus hampei and Anoplophora glabripennis*.

| GO ID | # Clusters | GO Description | GO Category | FDR |
|---|---|---|---|---|
| GO:0003964 | 17 | RNA-directed DNA polymerase activity | MF | 1.07738e-20 |
| GO:0046051 | 2 | UTP metabolic process | BP | 0.000447 |
| GO:0004518 | 5 | nuclease activity | MF | 0.000461 |
| GO:0045739 | 3 | positive regulation of DNA repair | BP | 0.000719 |
| GO:0007166 | 3 | cell surface receptor signalling pathway | BP | 0.000874 |
| GO:0006313 | 6 | transposition, DNA-mediated | BP | 0.000889 |
| GO:0015074 | 5 | DNA integration | BP | 0.001372 |
| GO:0031936 | 2 | negative regulation of chromatin silencing | BP | 0.001454 |
| GO:0042803 | 2 | protein homodimerization activity | MF | 0.002154 |

**Supplementary Table 6.** Extended gene model annotation statistics for the *Ips typographus* genome.

| | |
|---|---|
| Total sequence length | 236,816,287 |
| Number of genes | 23,923 |
| Number of exons | 121,796 |
| Number of introns | 97,873 |
| Number of CDS | 23,923 |
| Overlapping genes | 5,676 |
| Contained genes | 2,154 |
| Total gene length (bp) | 132,911,182 |
| Total exon length (bp) | 39,477,432 |
| Total intron length (bp) | 93,629,496 |
| Total CDS length (bp) | 29,963,361 |
| Shortest gene (bp) | 135 |
| Shortest exon (bp) | 1 |
| Shortest intron (bp) | 4 |
| Shortest CDS (bp) | 108 |
| Longest gene (bp) | 318,767 |
| Longest exon (bp) | 85,977 |
| Longest intron (bp) | 125,082 |
| Longest CDS (bp) | 75,285 |
| Mean gene length (bp) | 5,556 |
| Mean exon length (bp) | 324 |
| Mean intron length (bp) | 957 |
| Mean CDS length (bp) | 1,252 |
| Percent of genome covered by genes | 56 |
